# MATERNAL OBESITY MODULATES FOXO1 ACTIVATION AND ADIPOGENESIS IN NEONATAL MESENCHYMAL STEM CELLS

**DOI:** 10.1101/2025.08.01.668117

**Authors:** Sofía Bellalta, Theo Borghuis, Jelmer Prins, Marijke Faas, Torsten Plösch, Paola Casanello

## Abstract

Maternal obesity contributes to increase adiposity in the offspring. The fetal environment is crucial in determining the number of progenitor cells available for adipocyte formation and the extent of fat depots. In this context, adipogenesis is regulated by an intricate network of transcription factors, including FOXO1, which are responsive to external stimuli and ultimately result in the expression of the adipogenic marker PPARG. We hypothesized that the adipocyte progenitor cells, mesenchymal stem cells (MSCs), from neonates of women with obesity would exhibit a differential activation of the SIRT2-FOXO1-PPARG pathway, which is crucial in the regulation of early adipogenesis. In this study we isolated Wharton’s jelly-derived MSCs from neonates of women with obesity (BMI>30 kg/m², OB-MSCs) and women with normal-weight (BMI <25 kg/m², NW-MSCS) and induced them towards *in vitro* adipogenesis. OB-MSCs showed higher levels of FOXO1 and lower levels of acetyl-FOXO1 compared to NW-MSCs (p<0.05) and differential regulation of these proteins during early adipogenesis of OB-MSCs versus NW-MSCs (p<0.05). Further, acetyl-FOXO1 was higher in the cytoplasm as compared to NW-MSCs upon day 2 of adipogenesis (p<0.05). Finally, we found higher PPARG gene expression in OB-MSCs adipocytes compared with NW-MSCs adipocytes at day 21 of differentiation (p<0.05), denoting higher adipogenic potential. Our findings suggest that the maternal obesogenic environment influences the early modulation of FOXO1, which could lead to an increased adipogenic differentiation in MSCs from neonates of women with obesity, potentially increasing adipogenesis from their precursor pool. This could explain the higher adiposity in neonates of woman with obesity compared to normal-weight woman.

## Introduction

Maternal obesity is characterized by changes in circulating levels of nutrients, hormones, growth factors, cytokines and inflammatory mediators in the mother and the fetus [1, 2]. In this respect, new efforts are required to clarify how maternal obesity leverages an increased adiposity and obesity risk in the progeny [3, 4]. Recent evidence suggests that mesenchymal stem cells (MSCs), which are the embryonic adipocyte precursor cells, display an increased propensity towards adipogenic commitment when exposed to maternal obesity [5–7]. In this context, the mother’s excessive nutritional and inflammatory state could be transferred to the fetus by modulating and programming adipose tissue precursor cells.

The process of adipocyte differentiation encompasses several stages: multipotent mesenchymal precursors cells, preadipocytes, growth arrest, clonal expansion, terminal differentiation and mature adipocytes [8–9]. This process, known as adipogenesis, is governed by a complex network of transcription factors that regulate the expression of adipogenic genes. Forkhead box class O1 (FOXO1) emerges as a downstream target of the Insulin-induced kinase of phosphatidylinositol 3-kinase (PI3K) and protein kinase B (Akt) signaling pathway, which promotes growth in response to extracellular signals [10]. FOXO1 activity is important for MSC differentiation towards adipocytes, and it is regulated by post-translational modifications at multiple sites that determine nuclear export and import, and transcriptional activity [11].

FOXO1 participates in differentiation processes by regulating cell cycle, oxidative stress and lipid metabolism [12–14]. During early stages of adipogenic commitment, FOXO1 is induced to regulate growth arrest [15–16]. FOXO1 is also able to induce the expression of antioxidant enzymes important for protection of cells from the increasing Reactive Oxygen Species (ROS) during adipogenesis [17]. FOXO1 expression and activity undergo tight and temporal regulation during adipogenesis: while FOXO1 levels begin to rise in the early stage of differentiation, its transcriptional activity remains dormant until the clonal expansion phase concludes [15]. Depending on the specific environment and cellular distribution, FOXO1 can function as a transrepressor of the nuclear Peroxisome proliferator-activated receptor γ (PPARγ), which is the primary regulator of adipogenesis [18]. FOXO1 suppresses PPARγ through direct protein-protein interaction, while it activates PPARγ by releasing the binding [18]. Following the initial days of adipogenesis, FOXO1 is excluded from the nucleus. Consequently, FOXO1 cannot bind to repress PPARγ, thereby allowing PPARγ to induce adipogenic genes, which leads to the adipocyte commitment and maturation [14, 19, 20].

Sirtuins (SIRT) are NAD-dependent histone deacetylases which act as co-regulators in oxidative stress [21]. SIRT2 is the major sirtuin expressed in adipocytes, and it localizes in the cytoplasm [21]. Deacetylation of FOXO1 by SIRT2 is a regulatory mechanism during early adipogenesis. Jing *et al*. demonstrated that during early commitment of 3T3-L1 preadipocytes, overexpression of SIRT2 deacetylates FOXO1, leading to its nuclear translocation and repression of PPARγ [22]. On the contrary, knockdown of SIRT2 results in elevated FOXO1 acetylation, excluding FOXO1 from the nucleus [22]. This facilitates the adipocyte differentiation by reducing FOXO1 capacity to interact and suppress PPARγ transcription [22–24].

We have previously demonstrated that there are increased mitochondrial O_2_^•-^ levels and altered redox pathways in MSCs from the offspring of woman with maternal obesity [25]. Given the critical interplay between reactive oxygen species (ROS) and the SIRT2– FOXO1 axis during early adipogenesis [13, 22–24], we hypothesized that this pathway may also be dysregulated. In this study, we investigated the expression of FOXO1 and its regulation by SIRT2 and ROS during early adipogenic differentiation in MSCs from neonates born to mothers with normal weight versus those with obesity.

## Methods

### Experimental design

MSCs were isolated from the Wharton’s jelly of the umbilical cords of neonates from mothers with normal weight and obesity. The expression of SIRT2 and FOXO1 during early adipogenesis (day 0-5), and their response to ROS was assessed. The participation of FOXO1 was confirmed with the localization of acetyl-FOXO1. Further, adipogenesis was confirmed with PPARγ expression during adipogenesis and in final *in vitro* adipocytes (day 21).

### Subjects

Umbilical cords were anonymously taken from pregnant women with obesity or normal weight who delivered at the University Medical Centrum Groningen (UMCG) maternity ward, Groningen, The Netherlands, between July 2021, and July 2023. A pregestational maternal body mass index (BMI) >30 kg/m² was considered for the group of women with obesity (OB) and 18.5-24.5 kg/m² for normal-weight (NW) women. The exclusion criteria included women with gestational diabetes, preeclampsia, preterm birth, and neonatal complications [25]. All procedures were conducted according to the Helsinki Declaration.

### Isolation of Wharton’s jelly-derived MSCs

Umbilical cords were obtained from deliveries and directly processed to get MSCs primary cultures. MSCs were isolated through the explant method [25]. Briefly, the cord was cut and Wharton’s jelly explants were plated and cultured with Dulbecco’s modified Eagle medium (DMEM, Gibco, Thermo Fischer Scientific) (10 mM glucose), 10% fetal calf serum (FCS, Sigma-Aldrich) and 5000 UI/ml Penicillin-Streptomycin (Gibco, Thermo Fischer Scientific), maintained at 37°C in 5% CO₂. Subsequently, MSCs sprouted out of the explants and on day 10, cells were expanded into further passages. Unless otherwise stated, all experiments were performed in cells on passages 2-3. Cells were previously characterized according to the International Society of Cell Therapy criteria and stemness properties [25].

### Cell culture and adipogenic induction

Cells were seeded at 7.000 cells/cm² in 6-well plates and cultured with DMEM (Gibco, Thermo Fischer Scientific) (10 mM glucose), 10% FCS (Sigma-Aldrich), Penicillin-Streptomycin (Gibco, Thermo Fischer Scientific) and maintained at 37°C in 5% CO₂. When cells reached 80% confluency, adipogenesis was induced for 5 or 21 days with high glucose DMEM (25 mM glucose) supplemented with 10% FCS, 1% Penicillin-Streptomycin (Gibco, Thermo Fischer Scientific), 1 nM insulin (1Lonza Bioscience), 0.1 μM dexamethasone (Sigma-Aldrich) and 0.5 mM 3-isobutyl-1-methylxantine (Sigma-Aldrich) [26]. To simulate oxidative stress, during adipogenic induction, a mild oxidative challenge (250 μM hydrogen peroxide (H₂O₂) (Merck) was added to the culture every 48 hours. To inhibit SIRT2 expression, 10 μM AGK2 (Sigma-Aldrich) was added to the culture medium every 48 hours. This concentration was determined in pilot experiments (Supplementary figure S1).

### Protein collection and Western Blot analysis

MSCs were lysed with 100 μl RIPA buffer (Thermo Fischer Scientific) and 1 μl 100x Protease and Phosphatase Inhibitor Cocktail (Sigma-Aldrich) on day 0 (basal state) and days 1-5. Samples were sonicated and protein was quantified with a bicinchoninic acid (BCA) protein assay kit (Pierce, Thermo Fisher Scientific). 60 μg of proteins was run in 10% Sodium dodecyl-sulfate polyacrylamide gel electrophoresis (SDS-PAGE) for 1.5 hours at 120 mV, transferred to a nitrocellulose membrane (Bio-Rad) and blocked with 5% powdered non-fat milk (Friesland Campina) or BSA (Sigma-Aldrich) in 1x Tris-buffered saline plus 0.1% Tween T (TBST), for one hour. After blocking, the membranes were incubated overnight at 4°C with antibodies for FOXO1 (Invitrogen, 1:500), acetyl-FOXO1 Lys 294 (Invitrogen, 1:1000), SIRT2 (Cell Signaling, 1:500), PPARγ (Cell Signaling, 1:250) and β-actin (Cell Signaling, 1:10000) and α-tubulin (Proteintech, 1:1000). Thereafter, membranes were washed and incubated with secondary antibodies conjugated with HRP for one hour at room temperature (Rabbit Anti-Mouse and Goat Anti-Rabbit, Dako, Agilent, 1:5000). Only for experiments with ROS and AGK2, membranes were stained with Ponceau S for total protein as reference of protein quantity, because inhibition of SIRT2 can alter polymerization of actin [27]. Membranes were washed with TBST and signals were detected by chemiluminescence with SuperSignal West Pico PLUS Substrate (Thermo Fischer Scientific). Images were acquired with ChemiDoc XRS+ Imaging System (Bio-Rad). Data was quantified in ImageJ (NIH), and expressed relative to β-actin, α-tubulin or total protein expression.

### RNA isolation of MSCs on day 5 and day 21 of adipogenesis

Total RNA was isolated with TRIzol (Invitrogen) as described by the manufacturer [28]. Briefly, sample was homogenized, and layers were separated with chloroform (Merck). RNA was precipitated from the aqueous layer with 2-propanol (Merck). Pellet was reconstituted in water and RNA concentration and purity was measured with NanoDrop ND-100 UV-Vis spectrophotometer (NanoDrop Technologies).

### CDNA synthesis

A total of 500 ng of RNA were used for reverse transcriptase cDNA synthesis. The reverse transcription (RT) was performed with total RNA following standard protocol for RevertAid Reverse Transcriptase (Thermo Fischer Scientific Inc.). Briefly, samples were treated with DNAse I (Thermo Fischer Scientific Inc.) and incubated with Random Hexamer Primers, dNTPs set, Ribolock RNAse Inhibitor and RevertAid Reverse Transcriptase (Thermo Fischer Scientific) in a thermocycler (Biometra, Analytik Jena) following standard protocol.

### PPARG gene expression

Quantitative PCR (qPCR) was performed with FastStart Universal SYBR Green Master (Roche Diagnostics, Basel, Switzerland) for Peroxisome proliferator-activated receptor gamma (PPARG) gene according to manufacturer’s instructions. β-2-Microglobulin (B2M) and Glyceraldehyde 3-phosphate dehydrogenase (GAPDH) were used as housekeeping genes. qPCR was performed in Viia7 Real-time PCR System (Thermo Fischer Scientific). The threshold cycle (Ct) for gene amplification was detected and normalized as relative expression using 2^delta Ct (ΔCt = Ct [PPARG] – Ct [average housekeeping genes]) (Supplementary Table S1 – primers list).

### Localization of acetyl-FOXO1

Cells were seeded at 2.000 cells/cm² in Labtek chambers (Thermo Fischer Scientific). After 48 hours, in NW-MSCs and OB-MSCs adipogenesis was induced for 6 days as described above. At day 0, 2, 4 and 6 cells were washed with DPBS and fixed with PFA 2% (Sigma-Aldrich). After fixation, slides were washed with DPBS (Gibco, Thermo Fischer Scientific) and incubated with anti-acetyl-FOXO1 Lys 294 (Invitrogen, 1:500) for two hours. Further, cells were washed and incubated with secondary anti-rabbit Alexa Fluor 555 (Thermo Fischer Scientific, 1:500) for one hour. Cells were further stained with DAPI (Roche Diagnostics) for 15 minutes. Images were taken by Leica SP8 CLSM microscope (Leica Microsystems). For each image, Ilastik [29] was used to detect and separate nuclear-cytoplasmatic area. Further, Cell Profiler [30] was used to quantify mean fluorescent intensity of nuclear and cytoplasmatic area separately. Further, the nuclear/cytoplasmatic ratio was calculated. Experiments were done in duplicate for each subject.

### Supernatant SP3 digestion

Cells were cultured in the conditions described. After confluence, cells were starved from FBS for 24 hours. Later, the supernatant was collected and centrifuged at 400 x g for 10 minutes, and digestion of sample was performed based on the SP3 protocol previously described [31]. Briefly, samples were reduced with 10 μL dithiothreitol (10 mM) and proteins were bound to prewashed Cytiva SeraMag beads (Fisher Scientific). The samples were diluted in 50% acetonitrile and digested with 2.5 ng/μL trypsin (Promega) at 37 °C. Digestion was stopped by adding 10 μL 1% (v/v) formic acid, and the sample was prepared for the untargeted proteomics analyses on Evosep.

### Liquid Chromatography-Mass Spectrometry Analysis

Mass spectrometric analyses were conducted using an orbitrap mass spectrometer equipped with a nano-electrospray ion source (Orbitrap Exploris 480, Thermo Scientific) [31]. Briefly, peptide chromatographic separation was carried out on an Evosep system (Evosep One, Evosep). Ten percent of the digests were separated (30SPD workflow).

### Data analyses

LC-MS data were processed with Spectronaut (18.3.230830) (Biognosys) as described [31]. For data quantification of the supernatants, we used local normalization, and the Q-value filtering was set to the standard setting. Differentially expressed proteins were evaluated in STRING databases considering interaction with at least two out of the three proteins FOXO1, SIRT2 and PPARG [32].

### Statistical analysis

The parameters investigated were expressed as the median value and 25th and 75th percentiles (nonparametric distributions) in Graphpad Prism (Graphpad Inc). Mann– Whitney U test was used to compare between NW-MSCs and OB-MSCs. Paired Wilcoxon signed-rank test was used for paired data considering control and induced. Two-way ANOVA and Tukey’s post hoc test were used to assess the effects of obesity and time on PPARG gene expression, after transforming data to log2. P values < 0.05 were considered statistically significant.

## Results

### Basal FOXO1, acetyl-FOXO1 and SIRT2 protein expression in NW-MSCs and OB-MSCs

We evaluated protein expression for FOXO1, acetyl-FOXO1 and SIRT2 at day 0 (basal) in MSCs. We found that FOXO1 is significantly higher expressed in OB-MSCs compared to NW-MSCs (p = 0.03, Mann-Whitney U test; Figure 1). Levels of acetyl-FOXO1 and acetyl-FOXO1/total FOXO1 ratio were significantly decreased in OB-MSCs compared to NW-MSCs (p=0.05 and p=0.02, respectively, Mann-Whitney U test). SIRT2 was also higher expressed in OB-MSCs compared to NW-MSCs, although not significantly (p=0.08, Mann-Whitney U test).

**Figure 1.**
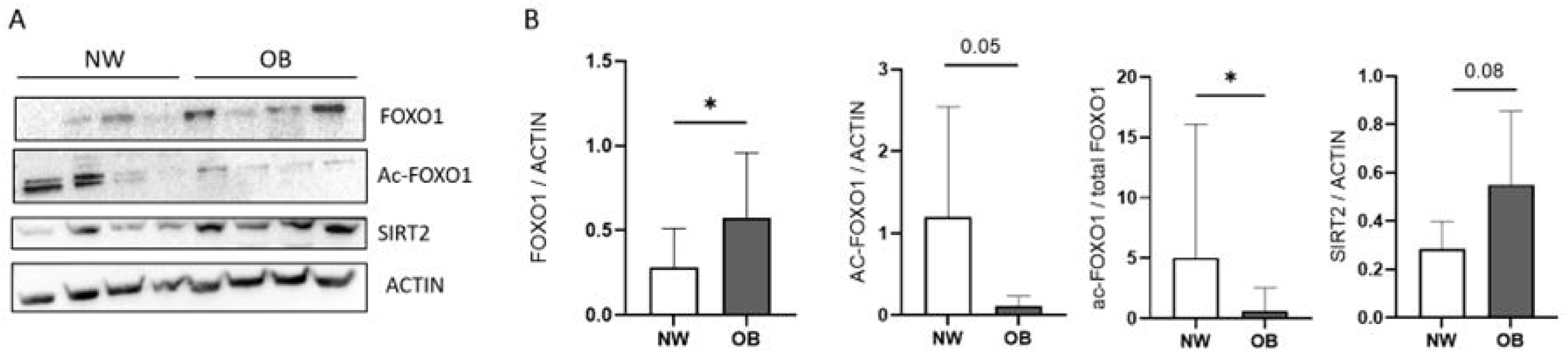
Expression of FOXO1, acetyl-FOXO1 and SIRT2 in NW-MSCs and OB-MSCs. **A**. Representative blot of expression of FOXO1, acetyl-FOXO1 and SIRT2 for both groups. **B.** FOXO1, acetyl-FOXO1, ratio of acetyl-FOXO1/total FOXO1 and SIRT2 on day 0. *p<0.05, Mann-Whitney U test (median ± range, n=8).

### Protein expression of FOXO1, acetyl-FOXO1 and SIRT2 in MSCs during early adipogenic differentiation

We next assessed the expression of these molecules in the first five days of adipogenic induction of MSCs. We found that FOXO1 was significantly increased from day 1 until day 5 versus day 0 in OB-MSCs (p<0.05, Paired Wilcoxon signed-rank test), while FOXO1 was only significantly increased on day 3 versus day 0 in NW-MSCs (p<0.05, Paired Wilcoxon signed-rank test, Figure 2). When we measured area under the curve (AUC) from day 0 until day 5, FOXO1 levels were significantly higher in OB-MSCs compared to NW-MSCs (p=0.04, Mann-Whitney U test, Figure 2). Interestingly, acetyl-FOXO1 significantly decreased at day 2 in NW-MSCs (p=0.04 for day 2 compared to day 0, Paired Wilcoxon signed-rank test) and not in OB-MSCs (p=0.07 for day 2 compared to day 0, Paired Wilcoxon signed-rank test). Further, levels of acetyl-FOXO1 are also significantly decreased at days 3, 4 and 5, compared to day 0 in NW-MSCs (p<0.05, Paired Wilcoxon signed-rank test), while acetyl-FOXO1 levels were only decreased on day 3 for OB-MSCs (p<0.05, Paired Wilcoxon signed-rank test), as compared with day 0. Total AUC of acetyl-FOXO1 was not different between both groups (Figure 2B). SIRT2 was significantly decreased on day 1 compared to day 0 in OB-MSCs (p=0.02, Paired Wilcoxon signed-rank test) but not in NW-MSCs (p=0.5, Paired Wilcoxon signed-rank test). There was no difference in SIRT2 levels over time for both groups. Total AUC of SIRT2 was not different between both groups (Figure 2C).

**Figure 2.**
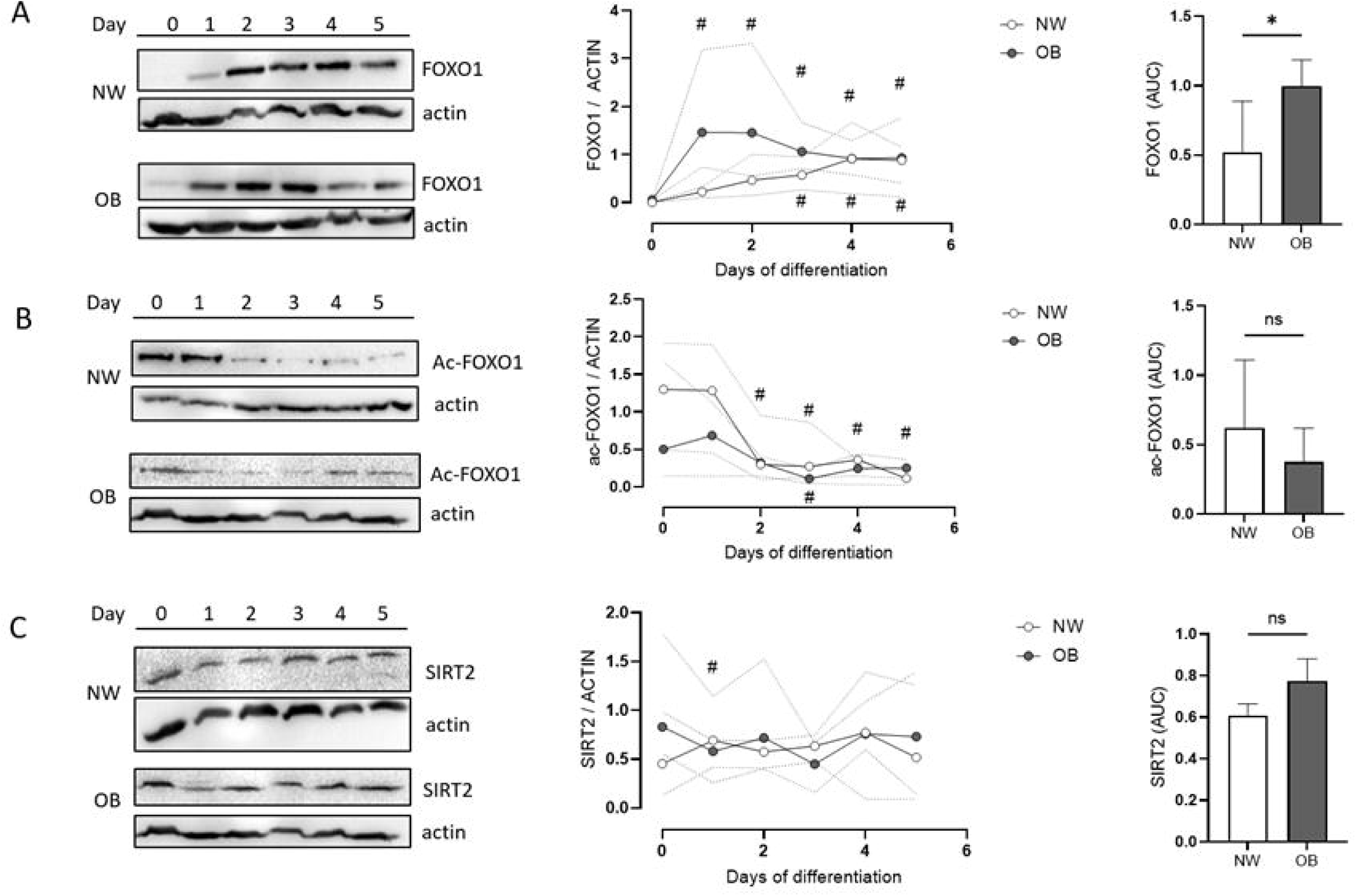
FOXO1 and SIRT2 levels during early adipogenesis in NW-MSCs and OB-MSCs. Cells were cultured with adipogenic induction media for 0, 1, 2, 3, 4 or 5 days and the expression of (A) FOXO1, (B) acetyl-FOXO1 and (C) SIRT2 was measured by western blot. Left: Representative western blots of NW-MSCs and OB-MSCs. Center: Levels of proteins over time in NW-MSCs and OB-MSCs. Right: Area under the curve (AUC) of levels of proteins in NW-MSCs vs OB-MSCs (median ± range, n=4). # p<0.05 Paired Wilcoxon signed-rank test compared to day 0 from the same group; *p<0.05 Mann-Whitney U test AUC.

### The effect of ROS on FOXO1, acetylation of FOXO1 and SIRT2 protein expression in NW-MSCs and OB-MSCs during early adipogenesis

In order to see whether the FOXO1-SIRT2 pathway is dependent on the presence of ROS during adipogenesis, we induced adipogenic commitment with and without the presence of H₂O₂ (250 uM) as a mild oxidative challenge. In accordance with the previous experiment, in NW-MSCs, FOXO1 is expressed at days 3 and 5 of adipogenesis. H₂O₂ significantly upregulates FOXO1 at day 3 (p = 0.01, Paired Wilcoxon signed-rank test), but not at day 5, compared to the untreated condition on the same day (Figure 3, top). Interestingly, for OB-MSCs (Figure 3, bottom), FOXO1 was also expressed at day 3 and 5 in the control condition, but there was no significant effect of H₂O₂ on FOXO1 expression on day 3 or day 5 of adipogenesis (p= 0.4 and p=0.2, Paired Wilcoxon signed-rank test). For acetyl-FOXO1, we found acetyl-FOXO1 at day 3 and 5 in the control condition, while our results show that H₂O₂ induced significantly higher levels of acetyl-FOXO1 at day 3 (p=0.01, Paired Wilcoxon signed-rank test), but significantly lower levels at day 5 (p = 0.01, Paired Wilcoxon signed-rank test) in NW-MSCs (Figure 3, top). However, we found that the effect of H₂O₂ is different in OB-MSCs: although acetyl-FOXO1 was found at day 3 and 5 in the control condition, at day 3 there was no response of acetyl-FOXO1 levels to H₂O₂ stimulation, whereas there is a tendency to an increased acetyl-FOXO1 levels at day 5 following H₂O₂ stimulation (p=0.06, Paired Wilcoxon signed-rank test, Figure 3B). SIRT2 levels were not affected by the presence of H₂O₂ during adipogenesis for NW-MSCs and OB-MSCs (Figure 3).

**Figure 3.**
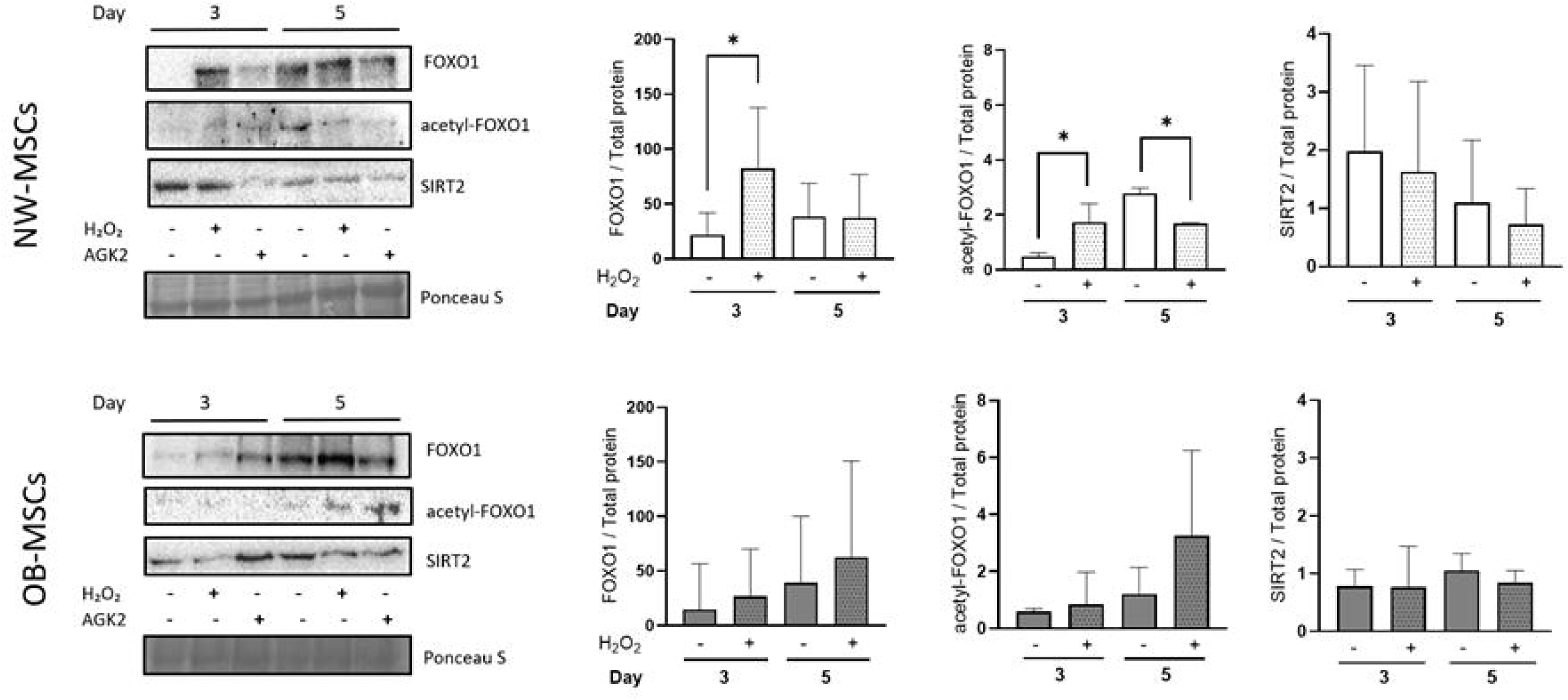
Induction of adipogenesis in NW-MSCs and OB-MSCs in the presence of H₂O₂. Cells were cultured with adipogenic induction media for 3 and 5 days, to assess the effect of H₂O₂ (250 μM) on FOXO1, acetyl-FOXO1 and SIRT2 levels. NW-MSCs (top), OB-MSCs (bottom) (median ± range, n=6). *p<0.05, Paired Wilcoxon signed-rank test. Representative western blots are shown in the left side.

### The effect of SIRT2 on FOXO1 and acetyl-FOXO1 in NW-MSCs and OB-MSCs during early adipogenesis

In Figure 2, we showed the presence of SIRT2 during early adipogenesis of MSCs (Figure 2). We next evaluated whether inhibition of SIRT2 affects the expression of FOXO1 and acetyl-FOXO1. For this, we inhibited SIRT2 with AGK2, a selective SIRT2 inhibitor [33] (Supplementary Figure S1). In NW-MSCs, treatment of the cells with AGK2 significantly inhibited SIRT2 protein expression at day 3 of adipogenesis (p = 0.04, Paired Wilcoxon signed-rank test), and non-significantly at day 5 (p=0.1, Figure 4, top). This effect was not seen in OB-MSCs (Figure 4, bottom). Further, there was no effect of AGK2 on FOXO1 expression in NW-MSCs and OB-MSCs (Figure 4 top and bottom). However, in NW-MSCs, AGK2 increased acetyl-FOXO1 on day 3, compared to the untreated condition (p = 0.03, Paired Wilcoxon signed-rank test, Figure 4, top), while at day 5, there was a tendency to lower levels of acetyl-FOXO1 in the presence of AGK2 (p = 0.06, Paired Wilcoxon signed-rank test, Figure 4, top). In OB-MSCs, acetyl-FOXO1 was higher at day 5 compared to the untreated condition (p=0.006, Paired Wilcoxon signed-rank test, Figure 4, bottom).

**Figure 4.**
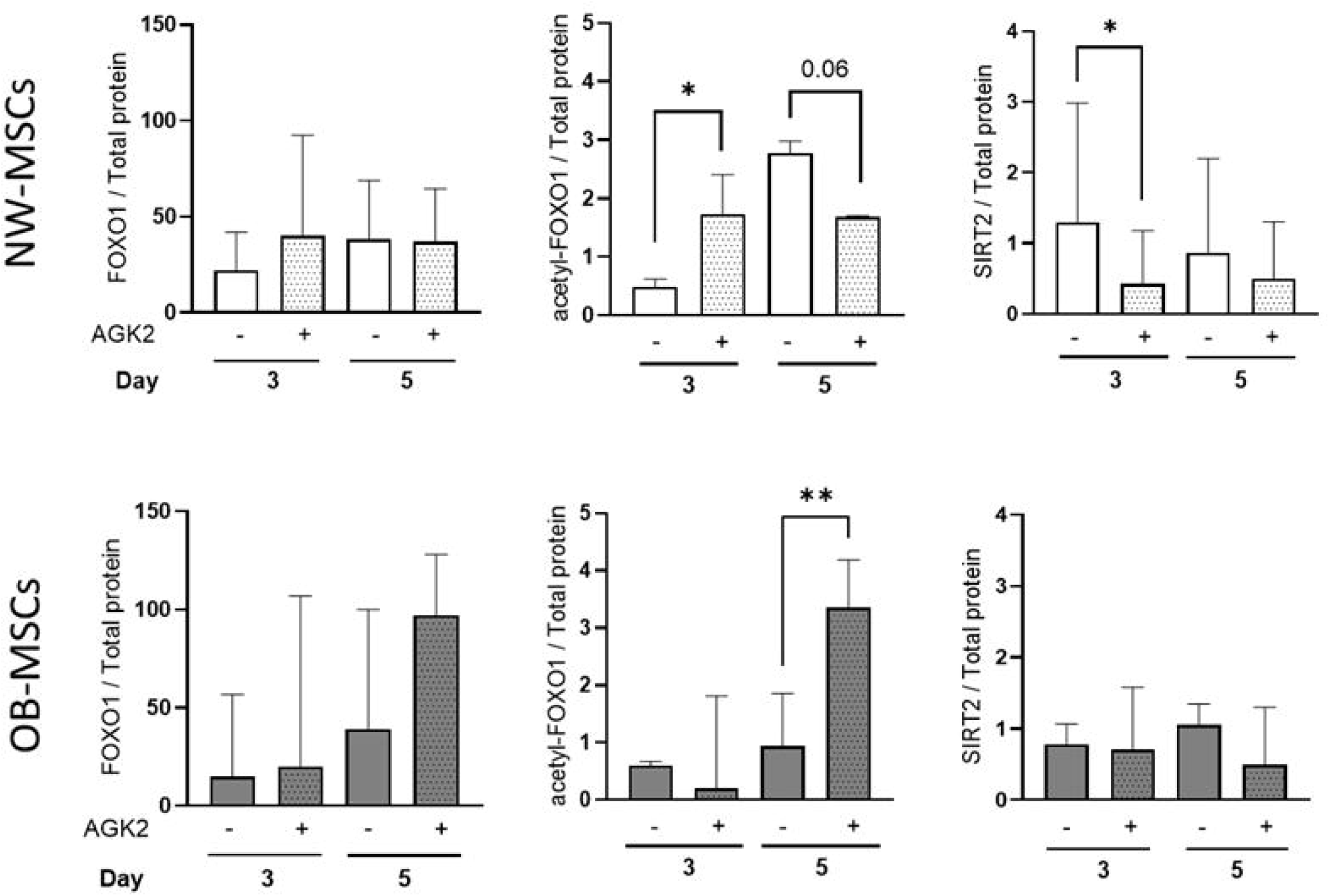
Induction of adipogenesis in NW-MSCs and OB-MSCs the presence of AGK2. Cells were cultured with adipogenic induction media for 3 and 5 days, to assess the effect of AGK2 (10 μM), a SIRT2 inhibitor, on FOXO1, acetyl-FOXO1 and SIRT2 levels. NW-MSCs (top), OB-MSCs (bottom) (Median ± range, n=6). *p<0.05, Paired Wilcoxon signed-rank test. Representative western blots are shown in Figure 3 (left side).

### Subcellular localization of acetyl-FOXO1 in OB-MSCs and NW-MSCs

Acetylation and deacetylation are post-translational modifications that regulate localization and trafficking of FOXO1 between nucleus and cytoplasm [34]. We assessed the subcellular localization of acetyl-FOXO1 at day 0, 2, 4 and 6 of adipogenesis, to evaluate its trafficking during early adipogenesis in both groups. At day 0, there was no difference in the nuclear/cytoplasmatic ratio of acetyl-FOXO1 between NW-MSCs and OB-MSCs, suggesting no difference in localization of acetyl-FOXO1 between NW-MSCs and OB-MSCs. Nevertheless, at day 2, we found that the nuclear/cytoplasmatic ratio of OB-MSCs was significantly lower as compared with this ratio of NW-MSCs (p=0.04, Mann Whitney U test, Figure 5). For day 4 and 6, there was no difference in the nuclear/cytoplasmatic ratio of acetyl-FOXO1 between both groups.

**Figure 5.**
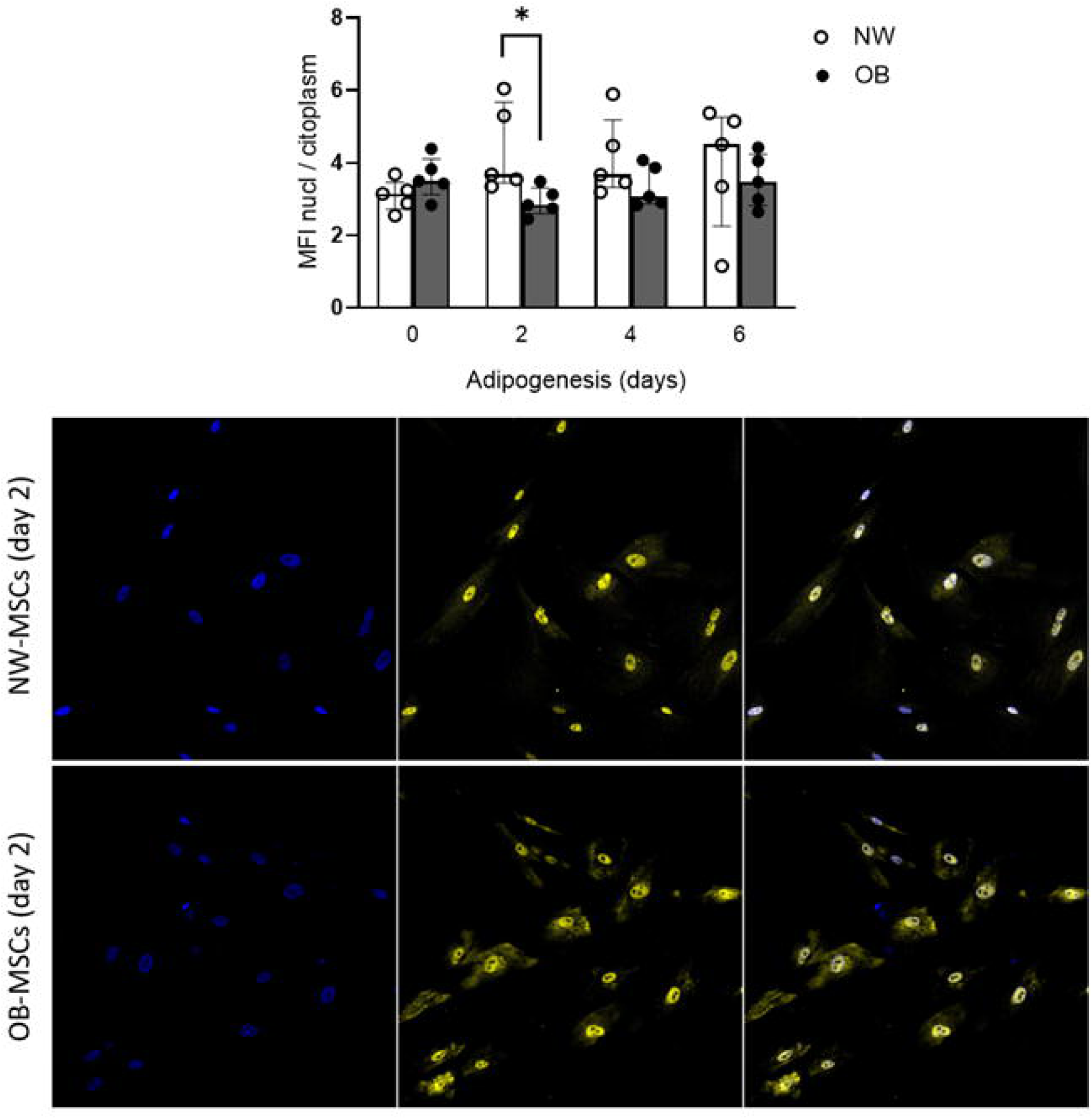
Subcellular localization of acetyl-FOXO1 during early adipogenesis. **A.** NW-MSCs and OB-MSCs were cultured with adipogenic induction medium and stained for acetyl-FOXO1 on days 0, 2, 4 and 6 of adipogenesis. Mean fluorescence intensity (MFI) of nuclear-cytoplasmatic ratio was calculated for localization. **B**. Representative images of day 2 for NW-MSCs (top) and OB-MSCs (bottom) (median ± range, n=6). *p<0.05, Mann-Whitney U test.

### The induction of PPARG gene and protein during adipogenesis in NW-MSCs and OB-MSCs

PPARG is known to regulate the adipogenic transcriptional machinery [35]. In our model, adipogenic commitment was confirmed with lipid droplets positive for Oil red O staining at day 21 (Figure 6A) and the presence of both isoforms, PPARγ1 and PPARγ2 (Supplementary Figure S2). Next, we aimed to assess whether PPARG gene expression was different between NW-MSCs and OB-MSCs, so we evaluated PPARG gene expression at day 5 (early adipogenesis) and day 21 (mature adipocyte). Our results show that there is an effect of the day of adipogenesis on the expression of PPARG (p=0.0001, two-way ANOVA), while there is no effect of obesity on the expression of PPARG (p=0.2, two-way ANOVA, Figure 6B). Post hoc testing showed an increased expression of PPARG on day 21 compared to day 5 of adipogenesis for both groups (p=0.001 for NW-MSCs and p=0.0001 for OB-MSCs, Tukey’s range test). On day 5 of adipogenesis, there were no differences in PPARG expression between NW-MSCs and OB-MSCs (p=0.7, Tukey’s range test). On day 21 of adipogenesis, OB-MSCs showed higher PPARG expression compared to NW-MSCs (p=0.03, Tukey’s range test, Figure 6B).

**Figure 6.**
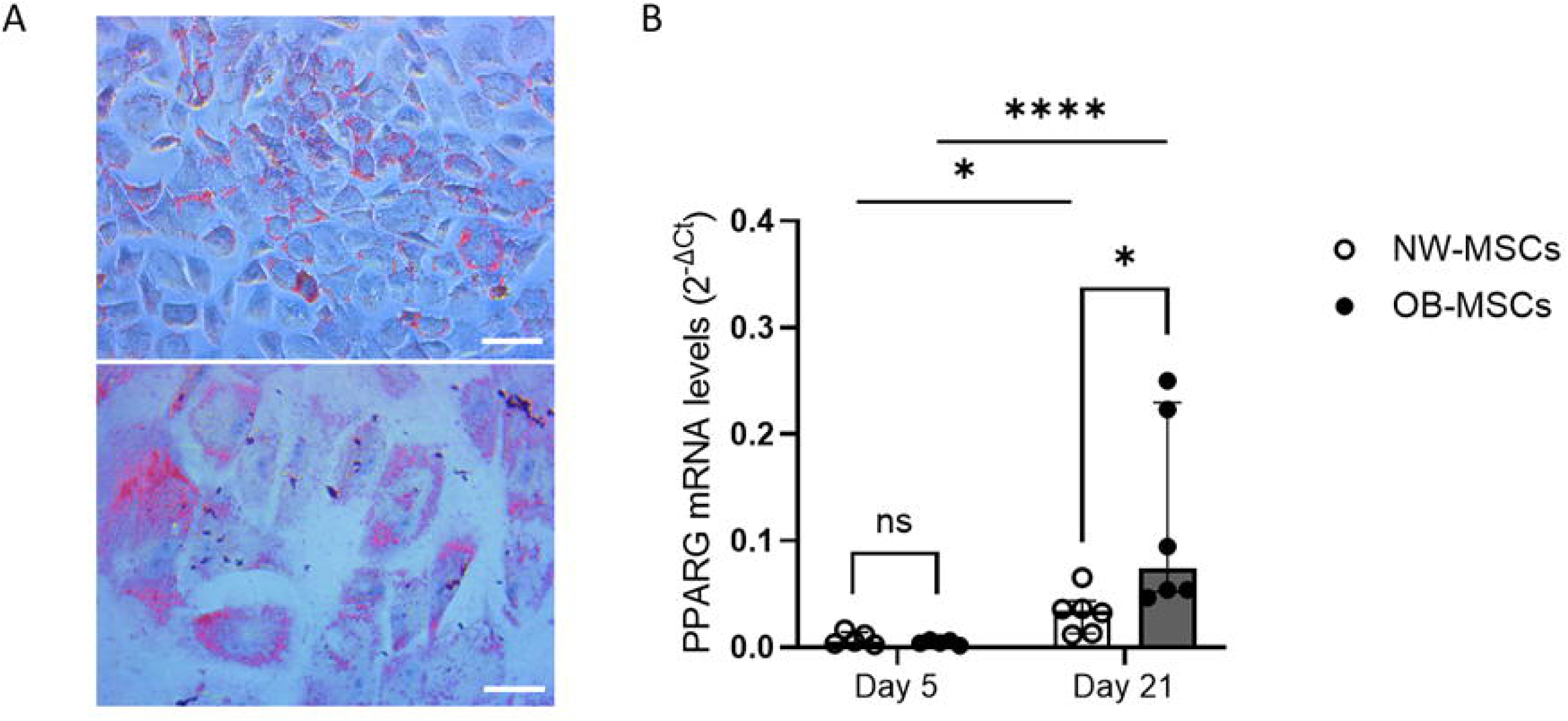
Adipogenic commitment of NW-MSCs and OB-MSCs. A. Lipid staining by Oil Red O in NW-MSCs on day 21 (scale bar: 100 μM top; 10 μM bottom). B. PPARG gene expression at day 5 and 21 of adipogenesis in NW-MSCs and OB-MSCs (Median ± interquartile range, n=6). Effect of day over PPARG (p<0.0001, two-way ANOVA); effect of obesity over PPARG (p=0.2, two-way ANOVA). *p<0.05 Tukey’s range post hoc test, ****p<0.0001 Tukey’s range post hoc test.

### SIRT2 and FOXO1 activity in the secretome of NW-MSCs and OB-MSCs

MSCs play an important role in the context of intracellular and intercellular communication via their paracrine activity [36, 37]. We evaluated the secretome profile of NW-MSCs and OB-MSCs in early adipogenesis (day 5) to see whether there were differences in secreted proteins that could be attributed to the FOXO1, SIRT2 and PPARγ pathways. Out of 660 differentially expressed proteins in OB-MSCs versus NW-MSCs, we identified 28 different proteins directly linked to SIRT2, FOXO1, PPARG, according to STRING databases (Table 1 and 2). From these 28 candidate proteins, the five proteins most abundant in the secretome of OB-MSCs versus NW-MSCs included nicotinamide phosphoribosyl transferase (NAMPT), caspase 3 (CASP3), GDH/6PGL endoplasmic bifunctional protein (H6PD), Cytochrome C (CYCS) and glucose-6-phosphate dehydrogenase (G6PD). The top five proteins less abundant in the secretome of OB-MSCs versus NW-MSCs included collagen alpha-1(I) chain (COL1A1), collagen alpha-2(I) chain (COL1A2), CD44 antigen, Peroxisome proliferator-activated receptor delta (PPARD) and Solute carrier family 2, facilitated glucose transporter member 1/GLUT 1 (SLC2A1). Table 1 and 2 summarizes the proteins with increased and decreased expression in the secretome of OB-MSCs versus NW-MSCs, respectively.

**Table 1.**
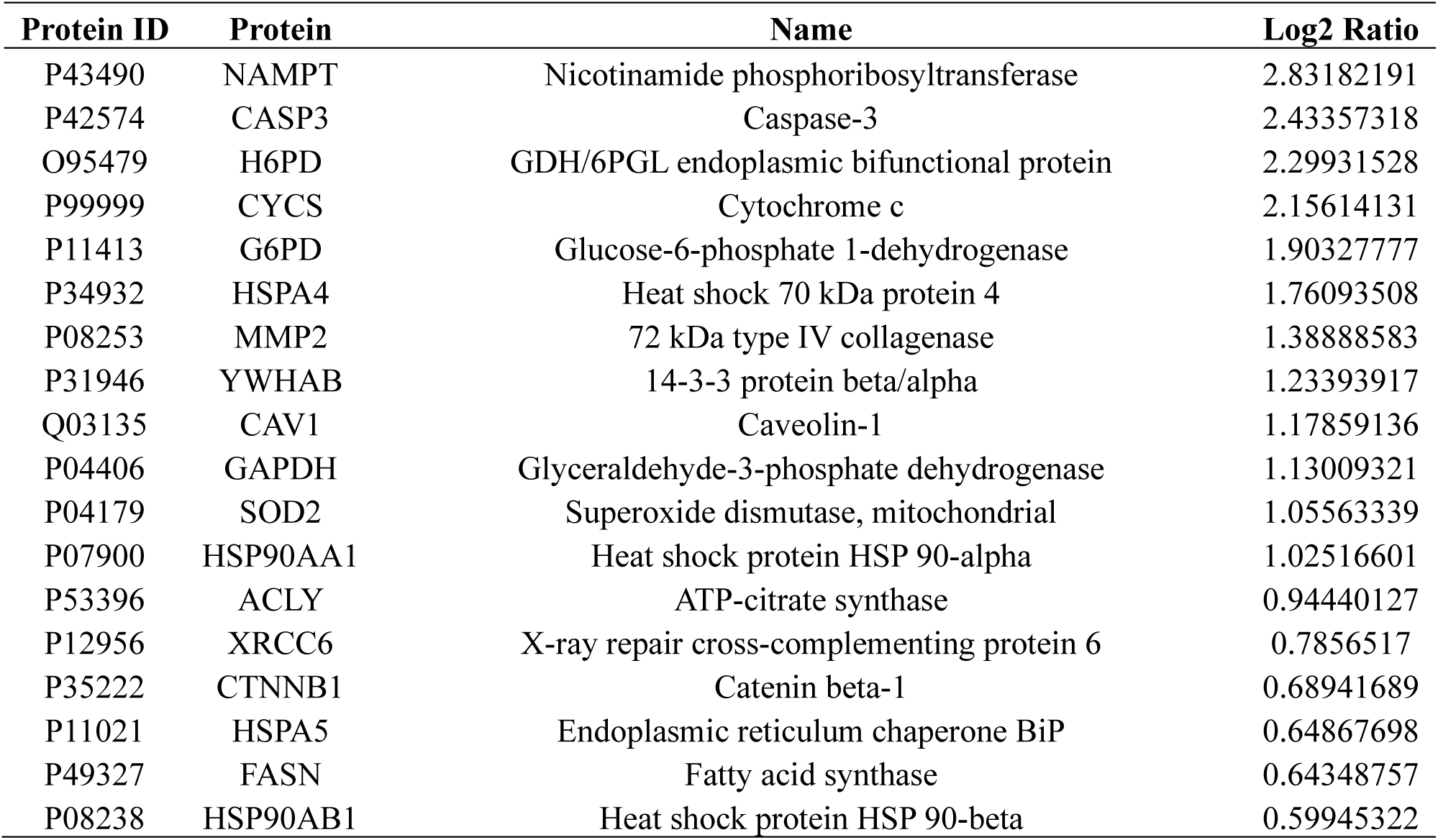
Increased proteins in the secretome from OB/NW-MSCs related to SIRT2, FOXO1 and PPARγ on day 5. STRING database.

**Table 2.** Decreased proteins in the secretome from OB/NW-MSCs related to SIRT2, FOXO1 and PPARγ on day 5. STRING database.

| Protein ID | Protein | Name | Log2 Ratio |
| --- | --- | --- | --- |
| Q92626 | PXDN | Peroxidasin homolog | -0.6686294 |
| P02649 | APOE | Apolipoprotein E | -0.7080032 |
| P02656 | APOC3 | Apolipoprotein C-III | -0.9501201 |
| P01137 | TGFB1 | Transforming growth factor beta-1 proprotein | -1.1204359 |
| P01023 | A2M | Alpha-2-macroglobulin | -1.1767215 |
| P11166 | SLC2A1 | Solute carrier family 2, facilitated glucose transporter member 1 | -1.2090146 |
| Q03181 | PPARD | Peroxisome proliferator-activated receptor delta | -1.312962 |
| P16070 | CD44 | CD44 antigen | -1.6520956 |
| P08123 | COL1A2 | Collagen alpha-2(I) chain | -1.7464086 |
| P02452 | COL1A1 | Collagen alpha-1(I) chain | -1.8029108 |

## Discussion

In this study we investigated the levels of SIRT2, FOXO1 and PPARγ in OB-MSCs and NW-MSCs and during early adipogenesis (day 0-5) and evaluated the effect of ROS on their levels during adipogenesis. We found that (1) basal and early adipogenic-induced FOXO1 is higher in OB-MSCs compared to NW-MSCs, which suggests a higher oxidative stress response and earlier regulation of cell cycle during adipogenesis; (2) acetyl-FOXO1 is lower in OB-MSCs, which means higher deacetylase activity and lower nuclear transport on day 0; (3) On day 2 of adipogenesis in OB-MSCs, acetyl-FOXO1 is higher in the cytoplasm as compared to NW-MSCs. This denotes less repression of PPARG and suggests its early activation towards adipogenic lineage commitment; (4) FOXO1 and acetyl-FOXO1 levels are affected in the presence of H₂O₂, therefore we propose that oxidative stress from maternal obesity could be modulating this early adipogenic pathway in neonates from mothers with obesity (Figure 7).

**Figure 7.**
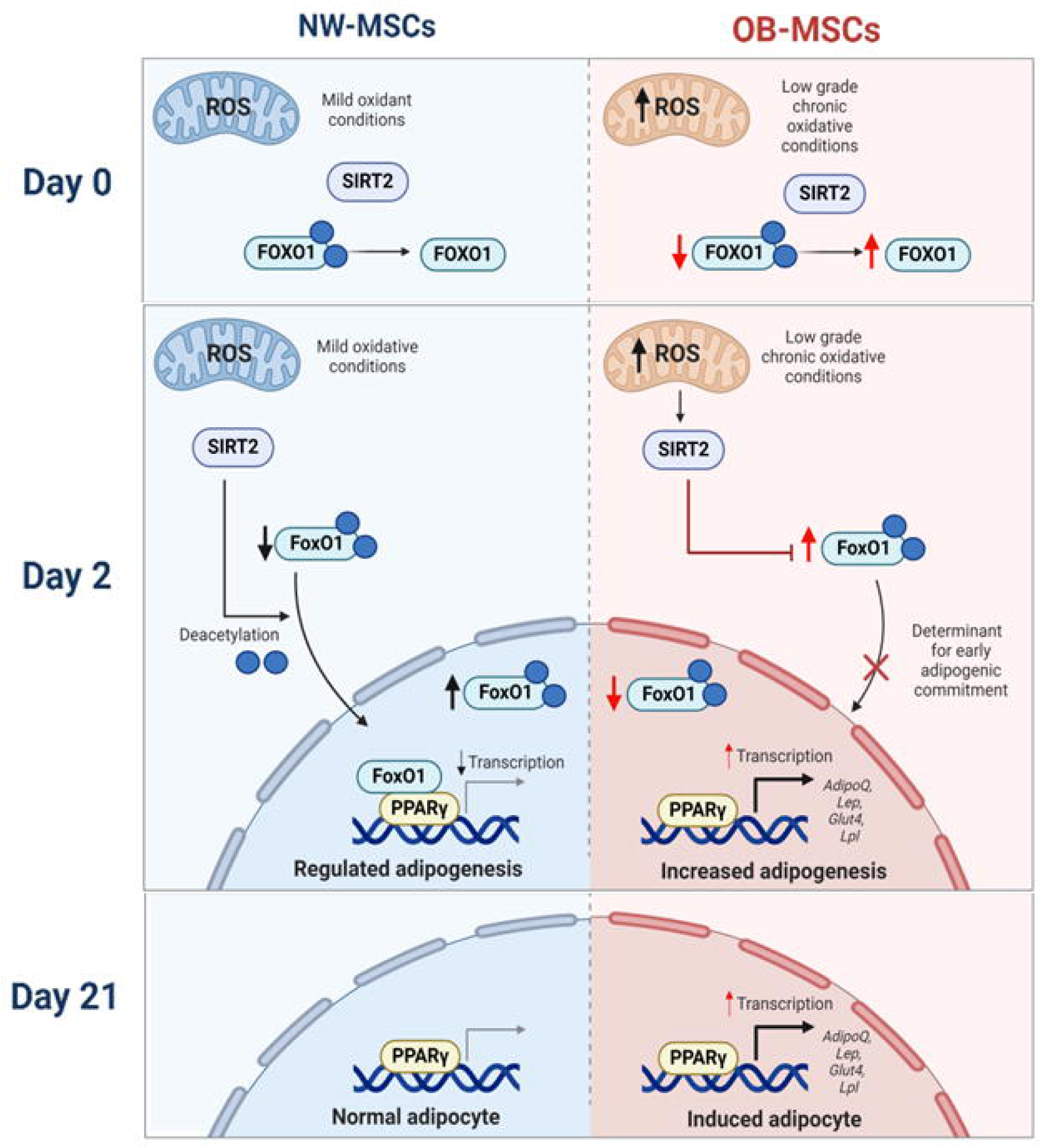
OB-MSCs show a higher FOXO1 and PPARG during adipogenesis. MSCs of neonates from mothers with obesity have higher levels of total FOXO1 and SIRT2, together with lower acetyl-FOXO1, which induces an early commitment in response to adipogenic induction. On early day 2 of adipogenesis, acetyl-FOXO1 localizes mainly in the cytoplasma for OB-MSCs, therefore releases PPARγ for adipogenic gene transcription. Finally, on day 21, *in vitro* adipocytes of OB-MSCs show higher levels of PPARG gene expression, compared to NW-MSCs. Created in BioRender.

FOXO1 is a transcription factor known to regulate cell cycle, oxidative stress and metabolism during adipogenesis [38]. In our study, we aimed to evaluate FOXO1 participation during adipogenic commitment in NW-MSCs versus OB-MSCs. We found that OB-MSCs have higher basal expression of total FOXO1 compared to NW-MSCs. It is known that FOXO1 responds to oxidative stress by binding to DNA to regulate antioxidant gene expression of superoxide dismutases, catalase, glutathione peroxidase, thioredoxins and peroxiredoxins [13, 17, 38–40]. Therefore, we propose that this higher expression of FOXO1 may be due to the exposition to oxidative stress during maternal obesity [25].

Our results also showed that OB-MSCs have higher levels and an earlier peak of FOXO1 induction compared to NW-MSCs during early adipogenesis (Figure 1 and Figure 2A), which suggests that OB-MSCs are inducing the adipogenic process earlier than NW-MSCs. These results are in agreement with the higher adipogenic commitment that has been previously reported in OB-MSCs [5–7]. FOXO1 is induced by adipogenic stimuli in early adipogenesis to regulate (1) cell cycle by stimulating growth arrest by the upregulation of p21, p27 to inhibit clonal expansion [41], (2) indirect activation of CEBP/s during adipogenesis [42] and (3) antioxidant enzymatic machinery because of physiological increase in ROS levels [13, 17, 38, 43]. We propose that due to the earlier expression of FOXO1, these three mechanisms are also induced earlier in the adipogenesis of OB-MSCs compared to NW-MSCs. Further studies are necessary to test this hypothesis.

Oxidative stress is known to regulate FOXO1 levels to activate antioxidant enzymes during adipogenesis [13, 17, 38–40]. In this study, H₂O₂ increased FOXO1 expression in NW-MSCs, but interestingly, FOXO1 was not responsive to H₂O₂ in OB-MSCs. FOXO1 has been described to regulate pathways widely associated with stress adaptation to confer resilience and ultimately promote lifespan [12, 13, 44]. Considering that we previously found oxidative stress in OB-MSCs and adaptive redox mechanisms [25], we suggest that the obesogenic intrauterine environment could play a role in the regulation of stress tolerance through FOXO1, and new studies should consider evaluating other FOXO1 mechanisms that could be affected by maternal obesity, such as gluconeogenesis, the insulin pathway and cell cycle regulation [13, 38].

FOXO1 activity is regulated by post-translational modifications, such as acetylation, that control DNA binding for transcription of target genes [14]. Our results showed lower levels of acetyl-FOXO1 and a lower acetyl-FOXO1/total FOXO1 ratio in OB-MSCs compared to NW-MSCs on day 0, suggesting that FOXO1 in OB-MSCs is mainly present in a deacetylated state and therefore could be actively binding to DNA for regulation of target genes such as cell cycle regulators and antioxidant enzymes [14, 15, 45, 46]. In both NW-MSCs and OB-MSCs, total cellular acetyl-FOXO1 levels decreased during adipogenesis and was significantly decreased from day 2 until day 5, compared to day 0. This denotes that decreased acetylation of FOXO1 is necessary during early days of adipogenic commitment and could implicate that deacetylated state of FOXO1 is active during adipogenesis [14]. The timing of the decreased acetyl-FOXO1 in MSCs is in accordance with previous findings showing that day 2 is known to be the timing for inducing growth arrest and activation of C/EBPs in 3T3-L1 preadipocytes during early adipogenesis [47].

Interestingly, we found that the protein 14-3-3/YWHAZ, which translocases deacetylated and phosphorylated FOXO1 between nucleus and cytoplasm, was upregulated at day 5 in OB-MSCs compared to NW-MSCs secretome (Table 1). This could be in line with the high levels of FOXO1 during early adipogenesis, since 14-3-3/YWHAZ is known to be secreted in cells with high activity of FOXO1 [48].

Acetylation and deacetylation of FOXO1 is not only vital for FOXO1 binding capacity to the DNA, but also for its translocation between the nucleus and cytoplasm [49, 50]. We found a lower nuclear/cytoplasmatic ratio of acetyl-FOXO1 in OB-MSCs compared to NW-MSCs at day 2. This represents a lower level of nuclear abundance and higher cytoplasmatic localization in OB-MSCs (results not shown), indicating that from day 0 to day 2 there may be trafficking from FOXO1 to the cytoplasm. These findings support the hypothesis that acetylation of FOXO1 in the cytoplasm contributes to its retention in the cytosolic compartment. As a result, FOXO1 is unable to repress expression of PPARγ. In this way, cytoplasmatic localization reduces the inhibitory effect of FOXO1 on adipogenesis, thereby facilitating differentiation in OB-MSCs towards adipocytes [15, 16, 18, 22, 24]. Altogether, this suggest that after day 2, which corresponds with the induction of cell cycle arrest, acetyl-FOXO1 levels decrease in both groups, marking the start of adipogenic differentiation. Moreover, since nuclear localization seems to be lower in OB-MSCs compared to NW-MSCs, there may be less repression of PPARγ in OB-MSCs as compared to NW-MSCs [18]. These data are in line with the suggestion that PPARγ becomes active for adipogenic commitment in OB-MSCs earlier than in NW-MSCs.

It is important to note that in our study we tested acetyl-FOXO1 as a mechanism for adipogenesis. As it has been found that acetylation increases phosphorylation of FOXO1 [45], further studies should also consider evaluating levels of phosphorylation of FOXO1. Additionally, we evaluated only acetylation at lysin 294, and it would be interesting to evaluate other acetylation sites that are important to regulate FOXO1 activity, such as lysine 245, 248 and 262 [12, 44, 46].

In agreement with the lower levels of acetyl-FOXO1 in OB-MSCs versus NW-MSCs in the basal state, we found that basal SIRT2 may be higher in OB-MSCs compared to NW-MSCs, although this is not significant. SIRT2 is a deacetylase and oxidoreductase regulated by NAD+/NADH levels [51]. SIRT2 is known to be higher expressed under mild oxidative stress where it has been linked to activation of superoxide dismutase as an antioxidant mechanism [52]. Therefore, we suggest that in OB-MSCs, like for FOXO1, SIRT2 is upregulated because of oxidative stress [25]. It is interesting to note that we also found higher levels of NAMPT in the secretome of OB-MSCs compared with NW-MSCs at day 5. NAMPT is the rate-limiting enzyme for NAD metabolism and hence for SIRT2 activity [53]. Another study reported lower levels of SIRT2 in MSCs and adipocytes from obese individuals [54]. Nevertheless, this study focused on MSCs from adipose tissue, together with mature adipocytes. The inconsistency with our results could be explained because these are MSCs from adults, who have been exposed to obesity and metabolic dysregulation much longer, while also many other metabolic pathways are affected [55].

Our results show that SIRT2 is not significantly changed by H₂O₂, however, there are changes in the levels of acetyl-FOXO1. This could indicate that H₂O₂ (250 uM) does not change SIRT2 protein expression but may affect its deacetylase activity. This may result in changes in acetyl-FOXO1, as seen in Figure 4. Unfortunately, we did not study the participation of other mechanisms regulating acetylation/deacetylation of FOXO1, which can be sensitive to H₂O₂ to attribute this effect [14]. This should be subject of future studies. SIRT2 was significantly inhibited by AGK2 in NW-MSCs, which is a specific inhibitor of SIRT2 [33] at day 3 of adipogenesis, and only slightly at day 5 of adipogenesis. However, there was no effect in OB-MSCs. This confirms our hypothesis of the robustness of SIRT2 in OB-MSCs, that could be associated to adaptation to oxidative stress [25]. Moreover, because acetyl-FOXO1 levels were increased on day 5 of adipogenesis in OB-MSCs in presence of AGK2, we suggest the participation of CBP/p300 acetylation or SIRT1 deacetylase activity in adipogenesis, and future studies should consider evaluating these other important mechanisms that regulate acetyl-FOXO1 in OB-MSCs [56, 57].

Our results suggest that the activities of SIRT2 and FOXO1 are regulated differently between NW-MSCs and OB-MSCs during early adipogenesis. With this, we hypothesized that PPARG gene expression would be different between both groups. Although we found lower nuclear localization of acetyl-FOXO1 in OB-MSCs compared to NW-MSCs on day 2, which strongly suggests less repression of FOXO1 on PPARG gene expression [18], we found no difference in PPARG gene expression on day 5 (early adipogenesis), which could indicate that other additional mechanisms inside the nucleus participate in the transcription of PPARG during that time period [14], and it would be interesting to measure direct binding of FOXO1-PPARG in NW-MSCs and OB-MSCs. Nevertheless, on day 21, PPARG expression was higher in OB-MSCs compared to NW-MSCs, which confirms a higher adipogenic commitment of this cells, as seen in other studies [5–7].

Not only intracellular pathways associated with differentiation towards adipocytes differ between NW-MSCs and OB-MSCs, also the secretome was different between both groups. On day 5 of adipogenesis, OB-MSCs showed increased and decreased expression of various secreted proteins related to SIRT2, FOXO1 and PPARγ, as compared to NW-MSCs. Interestingly, we found higher expression of lipogenic FASN and lower expression of PPARD and SLC2A1 in the secretome of OB-MSCs compared with NW-MSCs. FASN is a key enzyme for lipogenesis [58], PPARD is associated with lipid metabolism and oxidative phosphorylation in adipose tissue [59], while SLC2A1 (GLUT1) participates in glucose transport [60]. These findings suggest an altered metabolism in OB-MSCs compared to NW-MSCs. A different secretion pattern of proteins related to the SIRT2-FOXO1-PPARγ pathway in OB-MSCs versus NW-MSCs, suggest that the OB-MSCs may be communicating these changes to their microenvironment. As shown by the secretome data, the examination of candidate mechanisms through 14-3-3/YWHAZ, NAMPT, FASN, PPARD and SLC2A1 within whole cell lysates should be considered for subsequent studies.

In conclusion, in this study we found higher levels of FOXO1 and lower levels of acetyl-FOXO1 in OB-MSCs compared to NW-MSCs and differential regulation of these proteins during adipogenesis of OB-MSCs versus NW-MSCs. We also found higher PPARG gene expression in OB-MSCs adipocytes compared with NW-MSCs adipocytes at day 21 of differentiation. These findings suggest that the early modulation of FOXO1 can lead to an increased adipogenesis in MSCs from neonates of women with obesity, potentially increasing adipogenesis from their precursor pool.

