## Supplementary for "MATERNAL OBESITY MODULATES FOXO1 ACTIVATION AND ADIPOGENESIS IN NEONATAL MESENCHYMAL STEM CELLS"

**Supplementary Material**

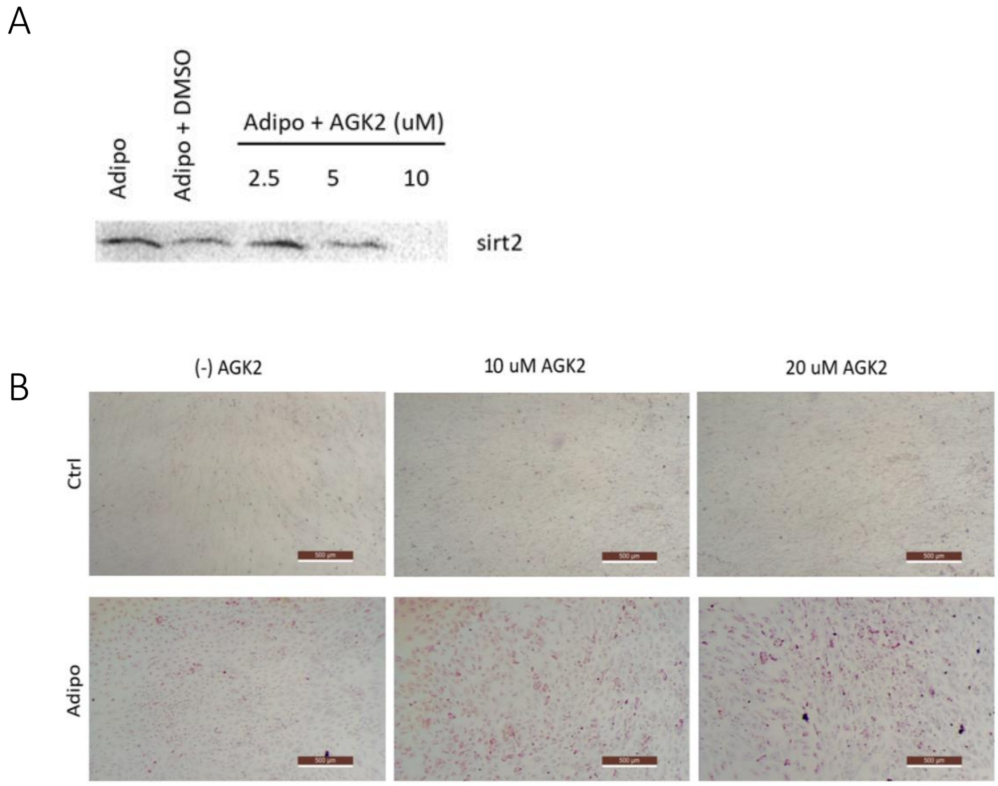


**Supplementary Figure S1. Inhibition of SIRT2 by AGK2 in NW-MSCs. NW-MSCs were cultured in the presence of AGK2 for inhibiton of SIRT2. A.** Protein expression of SIRT2 in NW-MSCs induced for 24 hours of adipogenesis in the presence of 2.5, 5 and 10 μM of AGK2, diluted in dimethyl sulfoxide (DMSO), indicating that 10 μM of AGK2 inhibits SIRT2 expression. **B**. NW-MSCs were induced for 21 days for adipogenesis in presence of 10 and 20 μM of AGK2. Lipid staining by Oil Red O (red) of NW-MSCs after 21 days of adipogenesis with 10 and 20 μM of AGK2, confirming higher adipogenesis by inhibiting SIRT2.

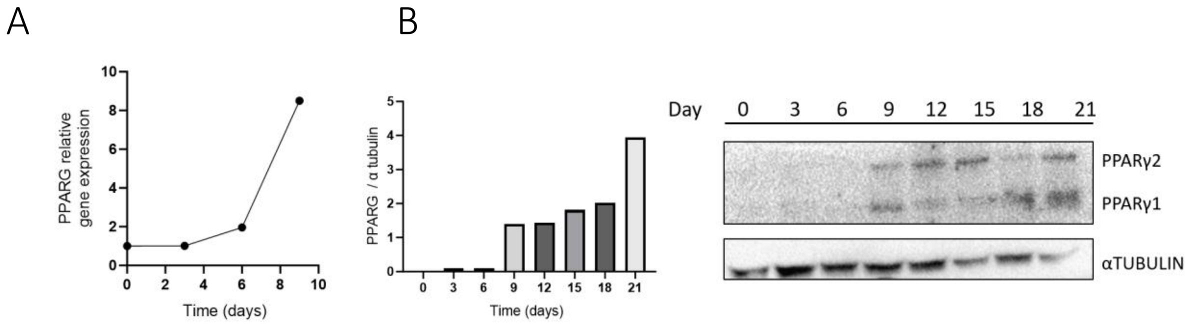


**Supplementary Figure S2. Gene and protein expression of PPARγ during adipogenesis of MSCs. A.** PPARG gene expression during 9 days of adipogenesis. **B**. PPARγ1 and PPARγ2 protein expression during 21 days of adipogenesis (n=1 NW-MSCs).

### Supplementary Table S1. Primers sequences and amplification conditions for mRNA RT-qPCR

| mRNA Target | Forward Sequence | Reverse Sequence | Efficiency (%) |
|---|---|---|---|
| *PPARG* | AACAGATCCAGTGGTTGCAG | AATTGCCATGAGGGAGTTGG | 94 |
| *GAPDH* | AGCCAACATCGTCAGACAC | GCCCAATACGACCAAATCC | 100 |
| *B2M* | GGTTTCATCCATCCATCCGACATT | ACGGCAGGCATACTCATCTT | 100 |
